# Heme-iron plays a key role in the regulation of the Ess/Type VII secretion system of *Staphylococcus aureus* RN6390

**DOI:** 10.1101/145433

**Authors:** M. Guillermina Casabona, Holger Kneuper, Daniela Alferes de Lima, Catriona P. Harkins, Martin Zoltner, Erik Hjerde, Matthew T.G. Holden, Tracy Palmer

## Abstract

The *Staphylococcus aureus* Type VII protein secretion system (T7SS) plays important roles in virulence and intra-species competition. Here we show that the T7SS in strain RN6390 is activated by supplementing the growth medium with hemoglobin, and its cofactor hemin (heme B). Transcript analysis and secretion assays suggest that activation by hemin occurs at a transcriptional and a post-translational level. Loss of T7 secretion activity by deletion of *essC* results in upregulation of genes required for iron acquisition. Taken together these findings suggest that the T7SS plays a role in iron homeostasis in at least some *S. aureus* strains.

## INTRODUCTION

Bacteria produce a number of different secretion machineries to transport proteins across their cell envelopes (1). Secreted proteins play essential roles in environmental adaptation and in pathogenic bacteria are frequently linked with the ability to cause disease. The type VII protein secretion system (T7SS) was discovered almost 15 years ago in pathogenic Mycobacteria. This system, also termed ESX-1, was shown to secrete two small proteinaceous T-cell antigens and to be essential for virulence (2–4).

Mycobacteria produce up to five different T7SSs (5, 6). In addition to ESX-1, ESX-5 also plays a key role in host interaction during pathogenesis (7). Of the other ESX systems in Mycobacteria, ESX-3 is the best-studied and is critical for siderophore-mediated acquisition of iron (8–10). Consistent with their diverse roles in the physiology and virulence of Mycobacteria, the ESX systems are differentially regulated. For example expression of ESX-1 is under indirect transcriptional control of the PhoPR two component system (11) that appears to respond to low pH, conditions that are found in phagolysosomes (12). ESX-5 expression is induced in response to phosphate starvation (13) whilst ESX-3 expression is de-repressed when cells are starved for iron or zinc (14, 15).

T7SSs are also found in other bacteria, in particular from the Gram positive low G+C firmicutes phylum (16). The similarity between the T7SS found in firmicutes and the well-studied Mycobacterial ESX T7SSs is limited, with the systems sharing only two common types of components. This has resulted in the T7SS of firmicutes such as *Staphylococcus aureus* being designated Ess or T7b to distinguish them from the better-characterised Mycobacterial T7a systems (17). One of the components shared between the two systems is a membrane-bound ATPase of the FtsK/SpoIIIE family termed EccC (T7a) or EssC (T7b). This component forms a hexameric assembly that likely acts as the motor protein and potentially also the translocation channel of the T7SS (18, 19). The second common component is at least one small protein of the WXG100 family, EsxA, which is secreted by the T7SS. In Mycobacteria, EsxA homologues are secreted as heterodimers with EsxB partner proteins (e.g. (20, 21) whereas in firmicutes EsxA is secreted as a homodimer (22).

The T7SS of *S. aureus* is encoded by the *ess* locus. In addition to EsxA and EssC, four further proteins encoded by the locus – EsaA, EssA, EssB and EsaB – are essential components of the secretion machinery (23–25). The *ess* locus is under complex transcriptional control by the alternative sigma factor σ^B^ and expression is also repressed by the two-component SaeSR system (26, 27). Experiments using mouse models of infection have indicated that the Ess system is required for virulence, in particular for the persistence of abscesses in the liver and kidney (23–25, 28). It is also required for colonisation and for intraspecies competition (25, 29). The secretion system appears to be highly expressed in mammalian hosts (30), and in at least one strain is transcriptionally activated by pulmonary surfactant (31). However, in laboratory growth media, although the secretion system components are produced, the machinery is poorly active and the levels of secreted substrates are relatively low (25, 29, 32).

In this study we have attempted to identify factors that activate secretion by the T7SS *in vitro*. We show that addition of hemin (heme B) enhances T7 secretion in at least two different *S. aureus* strains. Moreover, we also show that in the absence of a functional T7SS the laboratory strain RN6390 upregulates numerous genes involved in iron acquisition. Together our findings point to a novel role of the T7SS in *S. aureus* iron homeostasis.

## METHODS

### Bacterial strains and growth conditions for secretion assays

*S. aureus* strains used in this study are listed in Table 1. *S. aureus* strains were grown overnight at 37ºC with shaking in Tryptic Soy Broth (TSB). To test *S. aureus* growth with various media additives, strains were subcultured into either in TSB or RPMI (Sigma) as indicated, supplemented with the corresponding additives. Additives were made fresh and sterilised by filtration, and were dissolved in distilled water except for the hemin and other protoporhyrins which were dissolved in DMSO and hemoglobins which were dissolved in 0.1 M NaOH. For secretion assays, the indicated strains were grown to an OD_600_ of 2 and fractionated to give whole cell lysates and supernatant fractions as described previously (25). For growth curve analysis, additives were as follows: hemin 1, 2, 5 or 10 µM, 2,2’-birypidyl (Sigma) 200 μg ml^−1^.

**Table 1.** *S. aureus* strains used in this study.

| Strain | Relevant genotype or description | Source or reference |
| --- | --- | --- |
| RN6390 | NCTC8325 derivative, *rbsU*, *tcaR*, cured of $\phi 11$, $\phi 12$, $\phi 13$ | (51) |
| COL | Health-care acquired MRSA (HA-MRSA) | (52, 53) |
| USA300 | Community-acquired MRSA (CA-MRSA) | (54) |
| 10.1252.X | ST398-like isolate. Livestock associated | Roslin Institute,<br>Edinburgh |
| MRSA252 | HA-MRSA, representative of Epidemic MRSA-16 | (55) |
| HO 5096 0412 | HA-MRSA, representative of Epidemic MRSA-15 | (56) |

### Preparation of a polyclonal EsxB antibody

The EsxB (UniProt accession Q99WT7) coding sequence was PCR amplified from a synthetic gene (codon optimized for *Escherichia coli* K12 (Genscript)) using the forward primer 5’-GCGCGTCGACAATGGGCGGCTATAAAGGC-3’ and the reverse primer 5’-GCGCCTCGAGTTACGGGTTCACGCGATCCAGGC-3’, and cloned into the *SalI/XhoI* site of a modified pET27b vector (Novagen). The plasmid produces an N-terminal His_6_-tagged protein with a TEV (tobacco etch virus) protease cleavage site. The protein was expressed and purified as described previously (33), except in the final size exclusion chromatography step a HR 30/100 GL Superdex75 column (CV = 24ml, GE healthcare) equilibrated with 20 mM Tris pH 7.8, 100 mM NaCl was used. 2 mg purified EsxB (retaining a Gly-Ala-Ser-Thr sequence at the N-terminus after the cleavage step) was utilized as antigens to immunize rabbits for polyclonal antibody production in a standard three injections protocol (Seqlab, Goettingen, Germany).

### Western blotting

Samples were mixed with LDS loading buffer and boiled for 10 min prior to separation on bis-Tris gels. Western blotting was performed according to standard protocols with the following antibody dilutions: α-EsxA (25) 1:2500, α-EsxB 1:1000, α-EsxC (25) 1:2000, α-TrxA (34) 1:25 000. HRP-conjugated secondary antibodies (Bio-Rad) were used as per manufacturer’s instructions.

### RNA isolation and qPCR

For the RNA-seq analysis, three biological repeats of the indicated *S. aureus* strains were grown aerobically in TSB up to an OD_600_ of 1 at which point mRNA was prepared (in three technical replicates). Total mRNA was extracted, reverse transcribed and sequenced as described previously (35). The sequence reads from each individual dataset were mapped to the *S. aureus* NCTC 8325 genome using EDGE-pro (36), and quantification of transcript abundance and calculation of differential gene expression were performed using DEseq2 (37). DEseq2 use the Negative Binomial distribution as a model to compute p-values, and we regarded p > 0.05 as the probability of observing a transcript’s expression levels in different conditions by chance. Reads were aligned using the Tophat aligner (38) and to acquire a single transcriptome for each strain, the three assemblies produced by cufflinks were merged and the abundances of each sample were assembled using cuffquant. Differential expression was analysed using edgeR (39). Genes were considered to be differentially expressed when the logFC > 2 or < −2 and the q value < 0.05.

To isolate mRNA for RT-PCR, three biological repeats of the indicated *S. aureus* strains were grown aerobically in TSB in the presence or absence of 1**,i**M hemin up to an OD_600_ of 1 at which point mRNA was prepared. Total mRNA was extracted using the SV Total RNA Isolation Kit (Promega) with modifications as described in Kneuper *et al*. (25). Briefly, cell samples were stabilized in 5% phenol/95% ethanol on ice for at least 30 min and then centrifuged at 2770 *g* for 10 min. Cells were then resuspended in 100 µl of TE buffer containing 500 µg ml^−1^ lysostaphin and 50 µg ml^−1^ lysozyme and incubated at 37ºC for 30 min. Subsequently the manufacturer’s instructions were followed and the isolated RNA was subjected to a second DNase treatment using the DNA-free kit (Ambion). RNA was stored at −80ºC until use. To probe transcript levels, 500 ng of cDNA template was used with the following primer pairs: *esxA* (5’-TGGCAATGATTAAGATGAGTCC-3’ and 5,-TCTTGTTCTTGAACGGCATC- 3’ (25)), *esxC* (5’-AAGCATGCTGAAGAGATTGC-3’ and 5’-TCTTCACCCAACATTTCAAGC-3’) and 16S rRNA (5’-GTGCACATCTTGACGGTACCTA-3’ and 5’-CCACTGGTGTTCCTCCATATC-3’ (25)). Quantitative PCR was performed using a thermal cycler. Three technical replicates were prepared for each culture condition, using 2* Quantifast SYBR Green PCR master mix (Qiagen) according to manufacturer’s instructions. Standard curves were generated from serial 10-fold dilutions of genomic DNA. Amplification results were analysed with MxPro QPCR software (Stratagene) to give the levels of mRNA normalized to the level of 16S rRNA amplification in each sample. Results were further analysed in Microsoft Excel to calculate relative expression levels.

### Construction of an *esxA*-*yfp* transcriptional/translational fusion

The *yfp* gene was amplified without its start codon using Yfpfuse1 (3’ to 5’: GGAACTACTAGATCTTCAAAAGGC) and Xfpfuse2 (3’ to 5’: CCGGCGCTCAGAATTCTTATTTG) and cloned as a *Bgl*II/*Eco*RI fragment into pRMC2 (40), generating pRMC2-*yfp*. An approximately 500 bp region covering the *esxA* promoter, ribosome binding site and start codon was amplified using primers esxprom1 (3’ to 5’: GAATGGTACCGATTGTTGTTAAGATC) and esxprom2 (3’ to 5’: TTAGATCTTGCCATAACTAGAAACC) with RN6390 chromosomal DNA as template, and cloned as a *Kpn*I/*Bgl*II fragment into pRMC2-*yfp* to give plasmid pPesxA-yfp.

## RESULTS

### T7SS secretion in strain RN6390 is stimulated by supplementation with calcium ions, hemoglobin and hemin

Protein secretion systems are frequently activated in a post-translational manner, for example Type III secretion is activated by addition of the amphipathic dye Congo Red, or by calcium deprivation (41, 42) and the Type VI secretion system is activated by protein phosphorylation (43). We therefore sought to determine whether we could activate secretion by the T7SS in our standard laboratory strain of *S. aureus*, RN6390, by making empirical additions to the growth media. As shown in Fig 1A, panel (i), some secretion of the T7 core component, EsxA, could be detected when the strain was grown in either RPMI or TSB growth media. In general we noted that more EsxA was detected in the supernatant after growth in TSB than RPMI (Fig 1A).

**Figure 1.**
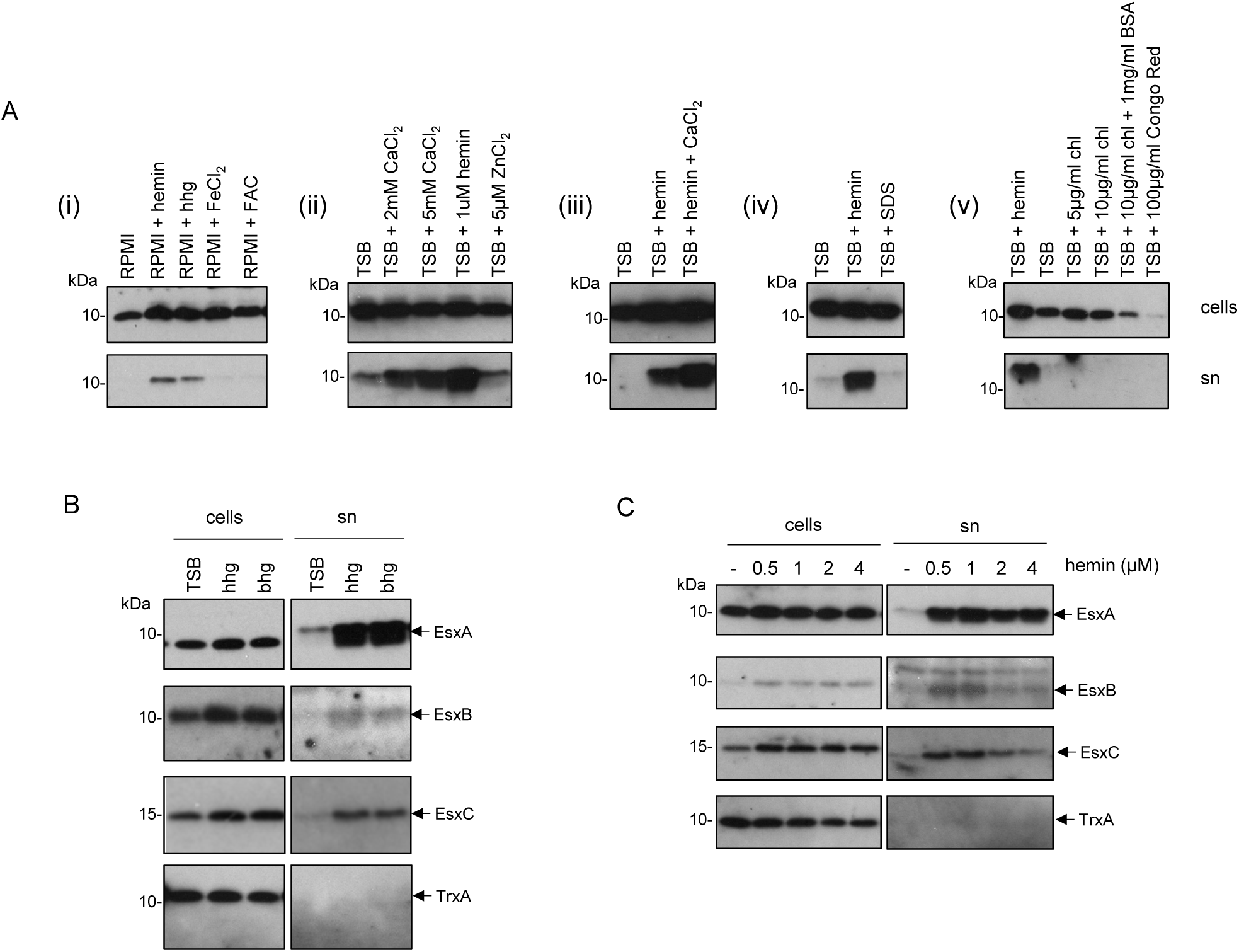
T7 secretion in strain RN6390 is stimulated by hemoglobin, hemin and mM concentrations of CaCl_2_. RN6390 was subcultured into either RPMI or TSB media, supplemented with the indicated additives, and grown aerobically until an OD_600_ of 2 was reached. Samples were fractionated to give cells and supernatant (sn), and supernatant proteins were precipitated using TCA. For each gel, 4 µl of OD_600_ 1 adjusted cells and 12 µl of culture supernatant were loaded. Final concentrations of additives, where they are not indicated on the figure are human hemoglobin (hhg) and bovine hemoglobin (bhg) 50µg/ml; hemin 1 µM; ferric ammonium citrate (FAC), FeCl_2_ 5 µM and CaCl_2_ 2mM. Chl – cholesterol, BSA – bovine serum albumin. (A) Western blots were probed with anti-EsxA antisera. (B and C) Western blots were probed with anti-EsxA, anti-EsxB, anti-EsxC or anti-TrxA (cytoplasmic control) antisera.

Both of these growth media lack an exogenously-added iron source, and RPMI is considered to be iron-limited (44). We therefore first tested the effect of exogenous iron sources on EsxA secretion. It can be seen that hemoglobin had a striking positive effect on EsxA levels in the culture supernatant for RN6390 grown both in RPMI and TSB media (Fig 1 A, panel i; Fig 1B). Fig 1B confirms that secreted levels of the T7 substrate proteins EsxB and EsxC (23, 24) were also similarly enhanced in the presence of hemoglobin. We tested whether other iron sources could also stimulate T7 secretion. Fig 1A (panel i) shows that neither ferric ammonium citrate (FAC) nor ferrous chloride stimulated EsxA secretion in RPMI, indicating that it was not a general effect of increased iron availability. The Mycobacterial ESX-3 T7SS is transcriptionally regulated by both iron and zinc (14, 15). However supplementation of the growth medium with 5 µM zinc was without detectable effect on EsxA secretion (Fig 1A, panel ii).

We next tested whether the iron-containing cofactor component of hemoglobin, hemin (heme B), could also enhance EsxA secretion. Fig 1A shows that supplementation of both RPMI (panel i) and TSB media (Fig 1A, panels ii-v) with 1µM hemin resulted in a marked increase in EsxA secretion. We confirmed that hemin had a similar stimulatory effect on the secretion of the T7 substrates EsxB and EsxC (Fig 1C). We conclude that hemoglobin and its cofactor, hemin, can positively regulate T7 secretion.

Next we tested whether calcium ions could also regulate secretion. Ca^2+^ is found at mM concentrations in mammalian blood and is also highly abundant in pulmonary surfactant (45). Fig 1A (panel ii) indicates that CaCl_2_ supplementation of TSB medium, at both 2mM and 5mM, increased the level of EsxA in the supernatant. Inclusion of 2mM CaCl_2_ alongside 1 µM hemin appeared to have additive effects over either supplement alone (Fig 1A, panels ii and iii). SDS, which has been shown to enhance *essC* mRNA levels (31) did not stimulate EsxA secretion (Fig 1A, panel iv). Finally, we tested whether either Congo Red or cholesterol, both of which stimulate protein translocation by Type III secretion system (41, 46) could increase EsxA secretion. However, Fig 1A (panel v) indicates that they did not enhance secretion of EsxA and moreover, Congo Red appeared to downregulate EsxA production.

Since hemin had the most marked enhancement on EsxA secretion, we examined whether increasing the concentration of added hemin would lead to further enhancement of secretion. Fig 1C shows that EsxA, EsxB and EsxC secretion were enhanced to similar levels in the presence of 0.5 and 1 µM hemin but at higher hemin concentrations secretion was reduced, and growth curves analysis indicated that higher hemin concentrations had a detrimental effect on growth of RN6390 (Fig S1A). In this context it has been noted previously that hemin concentrations of 5–10 µM are toxic to *S. aureus* (47).

Finally we undertook experiments to ascertain whether the empty (iron-free) protoporphyrin IX (PPIX) or zinc/copper –loaded PPIX could also stimulate EsxA secretion. Fig 2 shows that only hemin, the Fe-loaded form of PPIX, enhanced secretion of EsxA.

**Figure 2.**
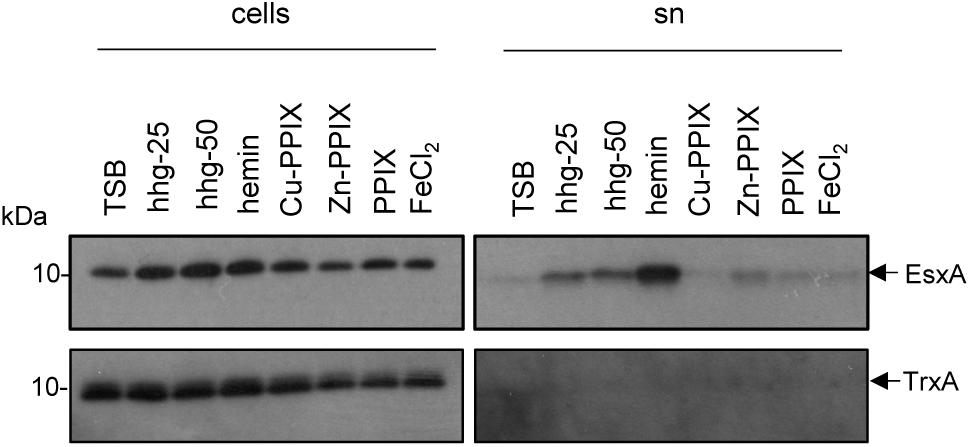
Stimulation of EsxA secretion is specific to the Fe-loaded protoporphyrin IX. RN6390 was subcultured into TSB medium supplemented with the indicated additives, and grown aerobically until an OD_600_ of 2 was reached. Samples were fractionated to give cells and supernatant (sn), and supernatant proteins were precipitated using TCA. For each gel, 4 µl of OD_600_ 1 adjusted cells and 12 µl of culture supernatant were loaded. Human haemoglobin (hhg) was added at either a final concentration of 25 µg ml^−1^ or 50 µg ml^−1^, FeCl_2_ at 5 µM and all other supplements at 2 µM. PPIX, protoporphyrin IX. Western blots were probed with anti-EsxA or anti-TrxA (cytoplasmic control) antisera.

### EsxA secretion is not induced by other oxidative stress compounds

In addition to acting as an iron source, at high concentrations hemin induces oxidative damage (48). To determine whether the hemin-induced hypersecretion of EsxA might be an oxidative stress response, we determined the effect of other oxidative stress agents on EsxA secretion. Fig 3A shows that in the presence of exogenous hydrogen peroxide there was potentially a small increase in the EsxA level in the supernatant. However in the presence of either diamide or methylviologen (paraquat) there was no stimulation of EsxA secretion and indeed the cellular level of EsxA appeared to be lower than the untreated sample. Co-supplementation of cultures with 1 µM hemin alongside diamide or methylviologen again resulted in hemin-dependent stimulation of EsxA secretion. We conclude that it is unlikely that the hemin induction of EsxA secretion in strain RN6390 is solely due to oxidative stress.

**Figure 3.**
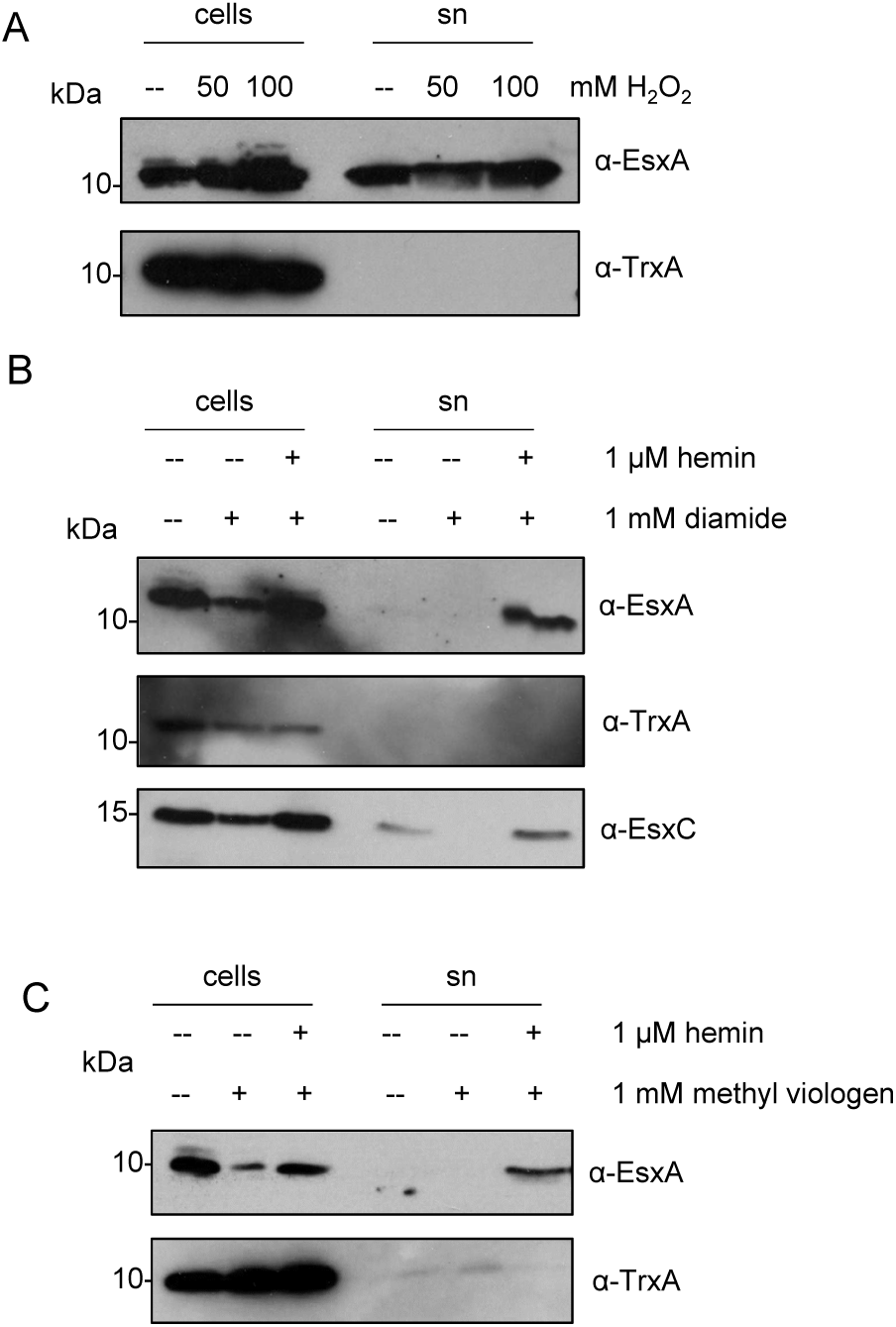
Effect of oxidative stress on EsxA secretion. *S. aureus* RN6390 was grown in the presence of the indicated concentrations of (A) H_2_O_2_, (B) diamide, or (C) methylviologen with or without the additional inclusion of hemin, and the secretion of EsxA was assessed by western blotting. For each gel, 5 µl of OD_600_ 1 adjusted cells and an equivalent of 15 µl of culture supernatant (sn) were loaded. Western blots were probed with anti-EsxA, anti-EsxC or anti-TrxA (cytoplasmic control) antisera.

### Hemin-induced hyper-secretion of EsxA is strain-dependent

Recent genomic analysis has revealed that there is genetic diversity at the *ess* locus across *S. aureus* strains. The *ess* loci were shown to fall into one of four different groupings, each of which is associated with a specific sequence variant of EssC, and with specific suites of candidate substrate proteins (35). We therefore undertook experiments to determine whether the hemin-induced stimulation of EsxA secretion was conserved across these groupings. COL is in the same EssC grouping as RN6390 (*essC1*) – both strains belong to the CC8 clonal complex, but are different sequence types (ST8 and ST250, respectively). COL has been noted previously to have a higher level of *in vitro* T7SS activity than RN6390 (25, 29). Fig 4 shows that EsxA secretion by COL is indeed higher than that of RN6390, and is comparable to the levels seen when RN6390 is grown with 1 µM hemin. Interestingly, hemin addition to cultures of COL grown in TSM medium had negligible effect on the level of EsxA secretion.

**Figure 4.**
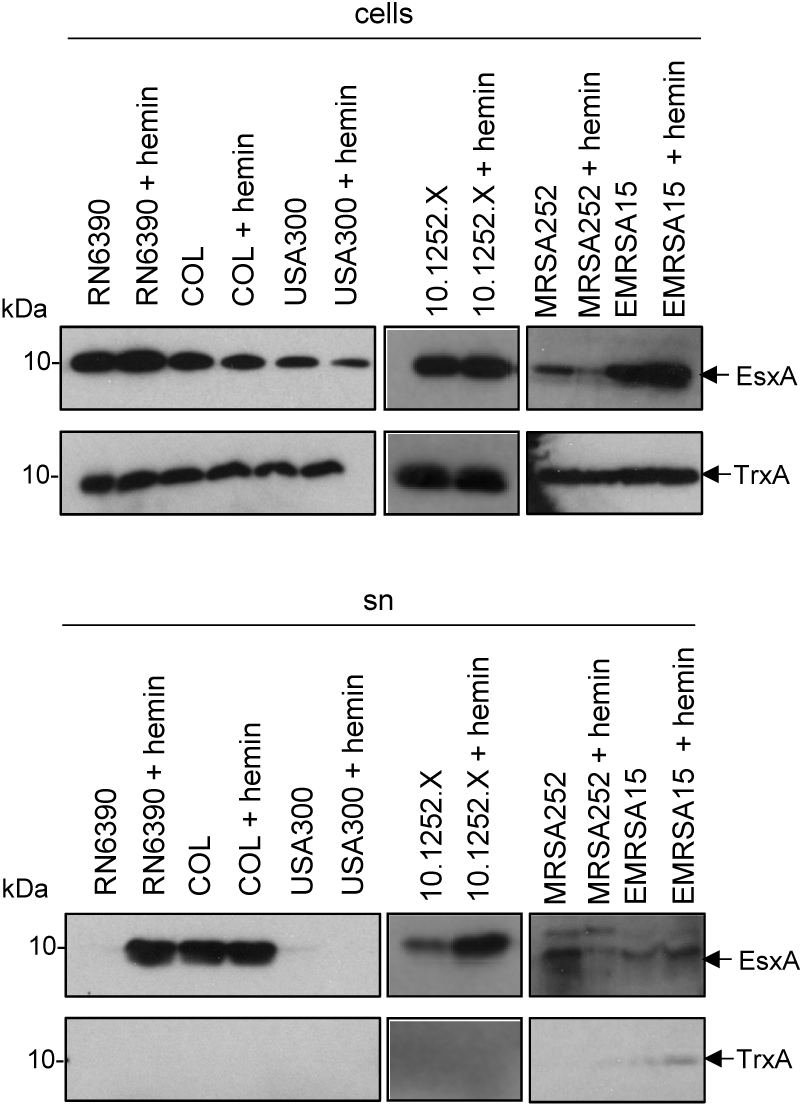
Hemin-induced stimulation of EsxA secretion in *S. aureus* is strain specific. The indicated *S. aureus* strains were subcultured into TSB medium, or TSB medium supplemented with 4 µM hemin, as indicated, and grown aerobically until an OD_600_ of 2 was reached. Samples were fractionated to give cells and supernatant (sn), and supernatant proteins were precipitated using TCA. For each gel, 4 µl of OD_600_ 1 adjusted cells and 12 µl of culture supernatant were loaded. Western blots were probed with anti-EsxA or anti-TrxA (cytoplasmic control) antisera.

We next examined the effect of hemin supplementation on an *essC2*-variant strain, *S. aureus* 10.1252.X. It can be seen (Fig 4) that 1 µM hemin also had a positive effect on EsxA secretion in this strain. By contrast, when strain MRSA252 (an *essC3* variant) was cultured with hemin, secretion of EsxA was reduced (and there also seemed to be less EsxA associated with the cellular fraction), suggesting a potential repression of *ess* expression in this strain (Fig 4). Finally when we examined the *essC4* strain variant, EMRSA15, there appeared to be a slight increase of EsxA levels in the supernatant in the presence of hemin, although we noted that there was some cell lysis in this strain as low levels of the cytoplasmic marker protein, TrxA, were also detected in the supernatant fraction. We conclude that the effect of hemin on EsxA secretion is strain-specific but that it clearly enhances secretion in two of the strains we examined.

### Hemin has a small transcriptional effect on *esxA* and *esxC* in RN6390

We next addressed the question whether hemin supplementation was increasing EsxA level in the supernatant due to transcriptional upregulation of the *ess* gene cluster. To this end, we isolated mRNA from RN6390 and the isogenic *essC* strain that had been cultured in TSB medium in the presence or absence of 1 µM hemin and used this to prepare cDNA. Since *esxA* is transcribed separately from the other 11 genes at the *ess* locus in RN6390, we undertook RT-qPCR with oligonucleotides designed to separately amplify *esxA* and *esxC*, normalising against 16s rRNA as an endogenous control. Fig 5A shows that there is a very small, but statistically significant, effect of hemin on both *esxA* and *esxC* transcription in the wild type RN6390 strain (1.5 – 2 fold). A similar small effect was also seen on the transcription of these genes in the presence of hemin when the T7SS was inactivated by deletion of *essC*. However, inspection of supernatant EsxA levels in the presence and absence of hemin (e.g. Fig 1A; 1C) indicates that there clearly a much greater than 2 fold increase in extracellular EsxA when hemin is added.

**Figure 5.**
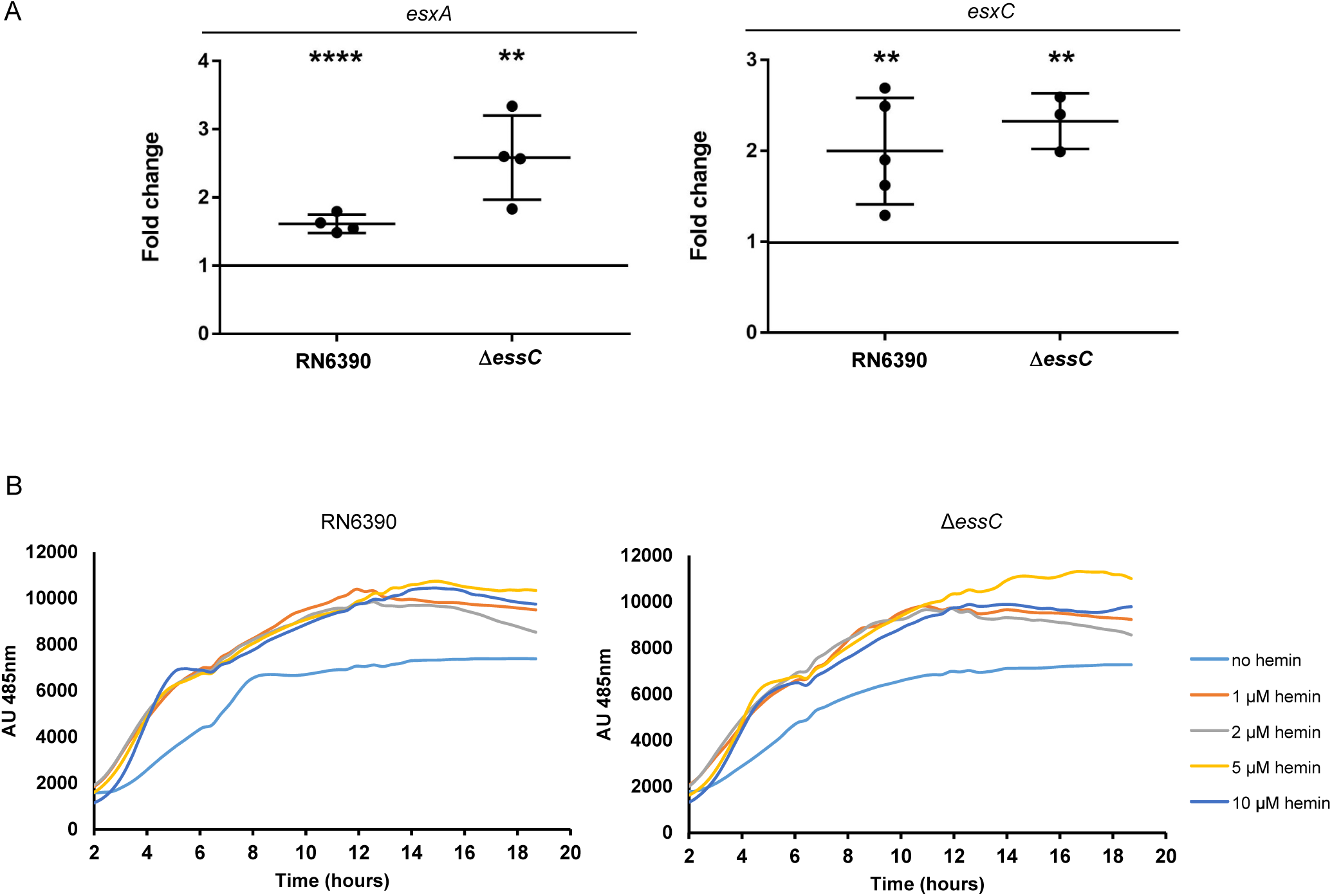
Hemin affects transcription of *esxA* and *esxC.* (A) *S. aureus* RN6390 and the isogenic *essC* deletion strain were grown aerobically in the presence or absence of 1 µM hemin to an OD_600_ of 1, at which point mRNA from at least three biological replicates was prepared as described in Methods. Relative transcription levels of the *esxA* and *esxC* genes were assayed by RT-qPCR (normalised against the level of 16S rRNA). *P* values are: **** < 0.0001; ** < 0.01. (B) *S. aureus* RN6390 and the isogenic *essC* deletion strain were cultured in the presence of the indicated concentrations of hemin for 18 hours in 96-well plates (200 µl volume) with shaking. YFP fluorescence was monitored at 485 nm and are measured in arbitrary units (AU) that were normalised to the growth at each time point.

To determine whether hemin may also affect translation of the *ess* genes, we constructed a plasmid-encoded fusion of the *esxA* promoter and ribosome binding site with *yfp* and monitored the fluorescence in the same two strains in the presence and absence of exogenous hemin. Fig 5B shows that there is a small (< 2-fold) effect of hemin supplementation on YFP fluorescence, consistent with the similar small effect seen on the *esxA* and *esxC* transcript levels seen by RT-qPCR. We conclude that hemin exerts both a transcriptional and most probably a post-translational effect on the T7SS in RN6390.

### Inactivation of RN6390 *essC* mounts an iron starvation transcriptional response

Taken together, the results so far indicate that heme iron has a striking effect on the secretion activity of the T7SS in *S. aureus* strain RN6390, potentially implicating the secretion system in iron homeostasis. To probe this further, we examined differences in the transcriptional profile between wild type reference strain RN6390 and an isogenic *essC* deletion mutant. Total RNA was prepared from exponentially growing cultures as described in Methods and RNA-Seq was used to investigate gene expression levels. As shown in Table 2 and Fig S2, a group of 41 genes displayed at least a log 2 fold statistically significant de-regulation in an *essC* mutant, with seven being down-regulated (excluding *essC* itself) and 34 genes being up-regulated. Interestingly, 25 of the up-regulated genes have known or implied roles in iron acquisition by *S. aureus* (49), these are listed in Table 3. Also included in Table 3 are the fold changes for all of the other genes involved in these iron acquisition pathways which did not reach our log 2 fold cut-off.

**Table 2.**
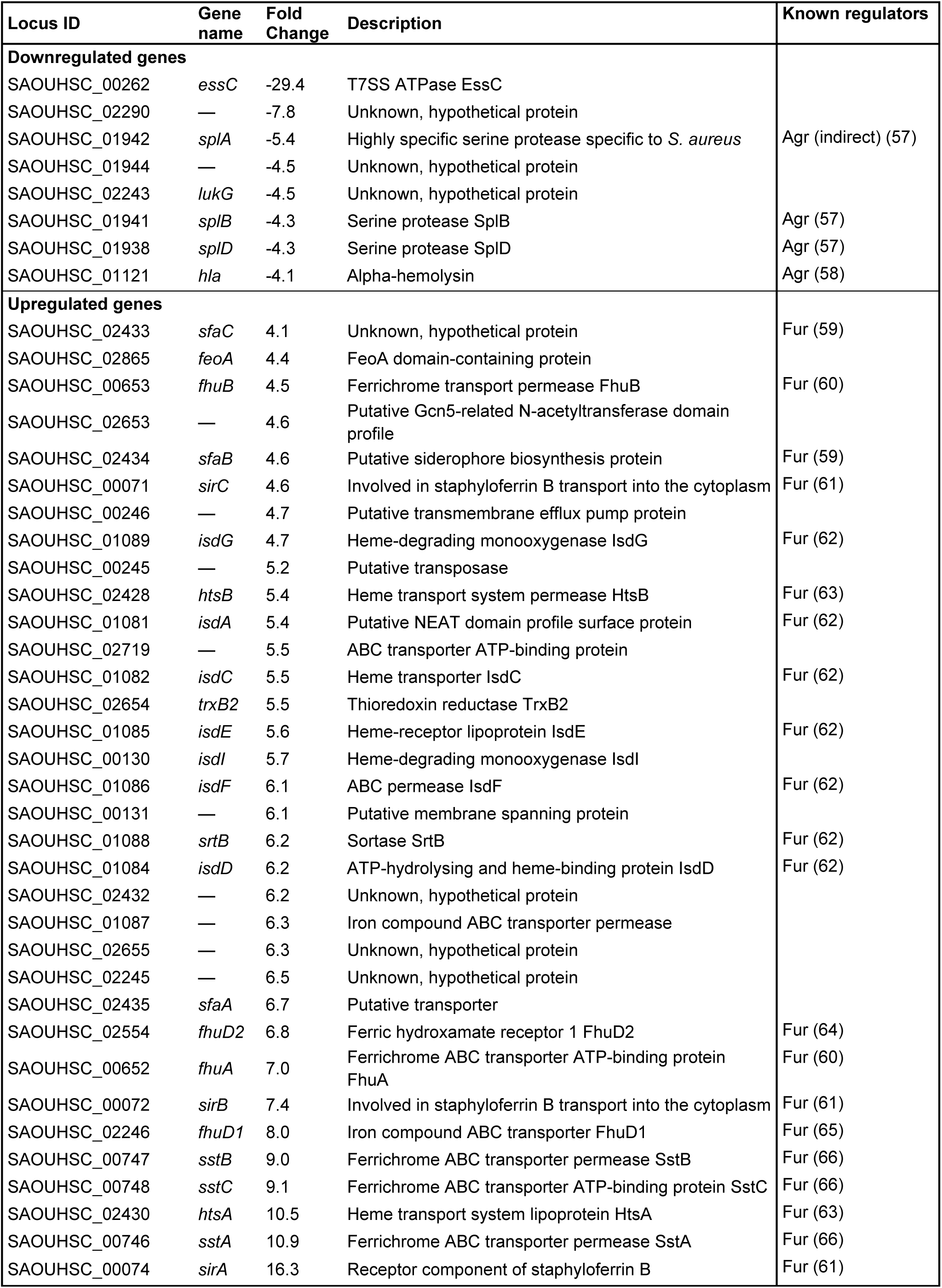
Genes differentially regulated (>log 2 fold) in the RN6390 *essC* deletion mutant, sorted by ascending fold change.

**Table 3.**
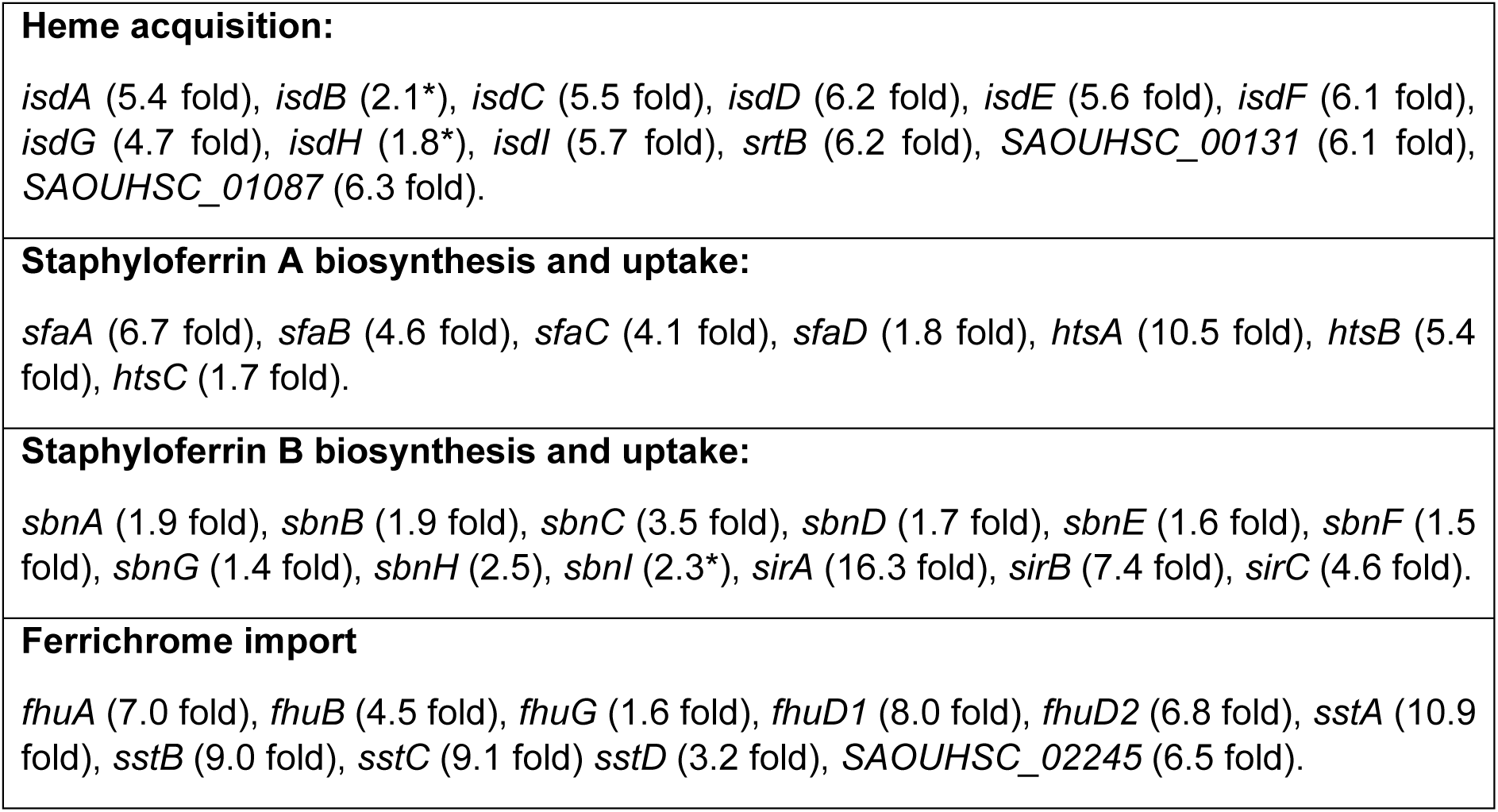
Genes involved in iron acquisition and level of upregulation in the *essC* mutant relative to wild type. *genes for which p value > 0.05 but were included for completeness.

It is apparent that many of the genes that encode the Isd machinery, which is involved in heme acquisition, are upregulated. Furthermore, genes for the biosynthesis and uptake of the two *S. aureus* siderophores, staphyloferrin A and staphyloferrin B, are also upregulated, as are genes encoding the Fhu machinery, which *S. aureus* uses to import xenosiderophores produced by other bacteria (49). All of these genes are known to be regulated by the ferric uptake regulatory (Fur) protein (Table 2). These findings indicate that inactivation of the T7SS by deletion of the *essC* gene prompts *S. aureus* RN6390 to mount an iron starvation response.

### The *S. aureus* RN6390 *essC* mutant is not iron starved but is more sensitive to hemin toxicity

Since the RNA-seq analysis suggested that the *essC* strain was iron starved, we next investigated the growth behaviour of strain RN6390 and the *essC* derivative under iron-limited growth conditions. Fig 6A shows that the addition of the iron chelator bipyridyl to TSB growth medium impaired the growth of the wild type strain, indicating that iron had now become a limiting factor. However, growth of the *essC* strain was almost indistinguishable from that of the parental strain in both the iron replete and iron limited TSB medium.

**Figure 6.**
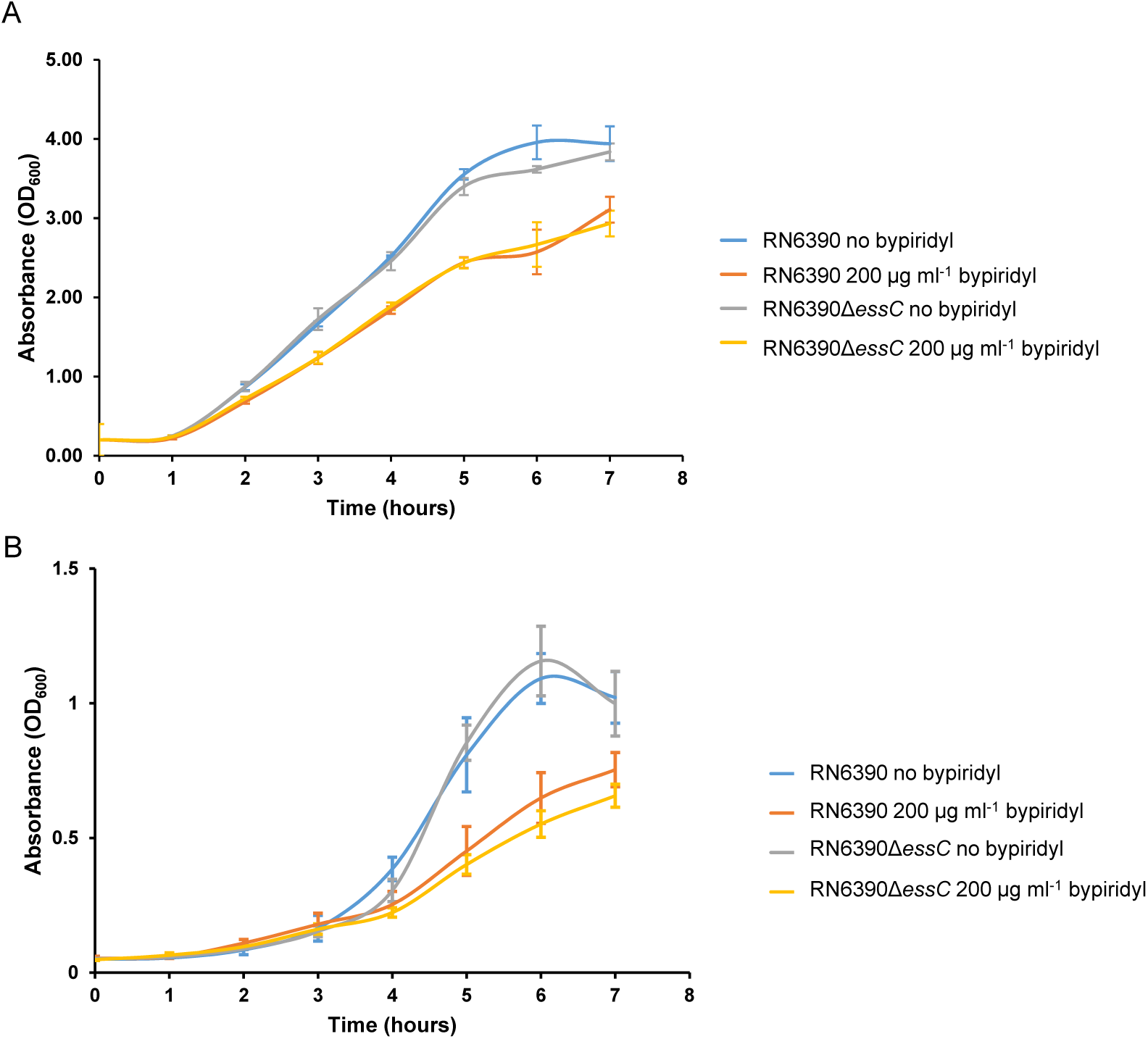
Effect of iron depletion on the growth of *S. aureus* RN6390. *S. aureus* RN6390 and the ∆*essC* isogenic mutant were grown in (A) TSB or (B) RPMI the presence or absence of 200 µg ml^−1^ 2,2’ bipyridyl at 37ºC with shaking and the OD_600_ was monitored hourly. Error bars are ± one standard deviation, *n*=3.

RPMI is a defined growth medium that is already iron deplete. Fig 6B shows that the parental strain grew more poorly in this medium than in TSB, and that additional inclusion of bipyridyl made little difference to the growth behaviour, consistent with the almost complete absence of iron in this defined medium. Again, the *essC* mutant strain showed essentially the same growth characteristics as the wild type. We conclude that inactivation of the T7SS does not lead to iron starvation.

Finally, we assessed whether the *essC* mutant was more sensitive to heme toxicity. Fig S1B indicates that at lower concentrations of hemin (1–2 µM) the *essC* mutant showed similar growth kinetics to the parental strain. However when the hemin concentration was increased to 5 µM or 10 µM the *essC* mutant clearly grew more slowly than the wild type. Taken together our findings demonstrate that the T7SS is upregulated in response to hemin in strain RN6390 and that this may subsequently contribute to the protection of *S. aureus* from hemin-induced toxicity.

## DISCUSSION

In this work we have sought to identify conditions that stimulate T7 protein secretion in our model strain of *S. aureus*, RN6390. After screening a range of media additives we identified hemoglobin and its cofactor hemin (heme B) as a secretion activator. Using variants of hemin that were lacking bound iron or that were loaded with alternative metals indicated that it was specifically the iron replete form of hemin that was required to stimulate secretion. Analysis of transcript levels of two genes encoded at the T7/*ess* gene cluster demonstrated only a small (approximately 2-fold) effect of hemin on *ess* gene transcription and suggested that the regulation of T7 activity is likely to be largely post-translational. How the T7SS is regulated at the post-translational level has not yet been established.

A further link between the T7SS and iron homeostasis was also identified by comparative RNA-seq analysis between the RN6390 wild type strain and a strain lacking the essential T7 secretion gene, *essC*. It was seen that loss of T7 activity was associated with upregulation of a large group of Fur-regulated genes that are required for iron acquisition. However, despite this, the *essC* mutant strain was not phenotypically iron starved. Fig 7 presents a speculative model that could account for these findings. According to the model, the T7SS in strain RN6390 secretes one or more substrate proteins involved in iron/heme binding and homeostasis (Fig 7A). It should be noted that the ESX-3 T7SS from both *Mycobacterium tuberculosis* and *Mycobacterium smegmatis* plays a key role in iron homeostasis, under iron replete and iron sufficient conditions, and secretes at least two substrates involved in siderophore-mediated iron uptake (10, 50). Inactivation of T7 secretion by loss of the core component EssC results in mislocalisation of the iron-binding substrate protein/s to the cytoplasm (Fig 7B). These cytoplasmic substrate/s titrate iron atoms away from the iron-binding transcriptional regulator Fur, resulting in the observed activation of Fur target genes. The identity of candidate T7 substrate proteins involved in iron homeostasis will be the subject of future analysis.

**Figure 7.**
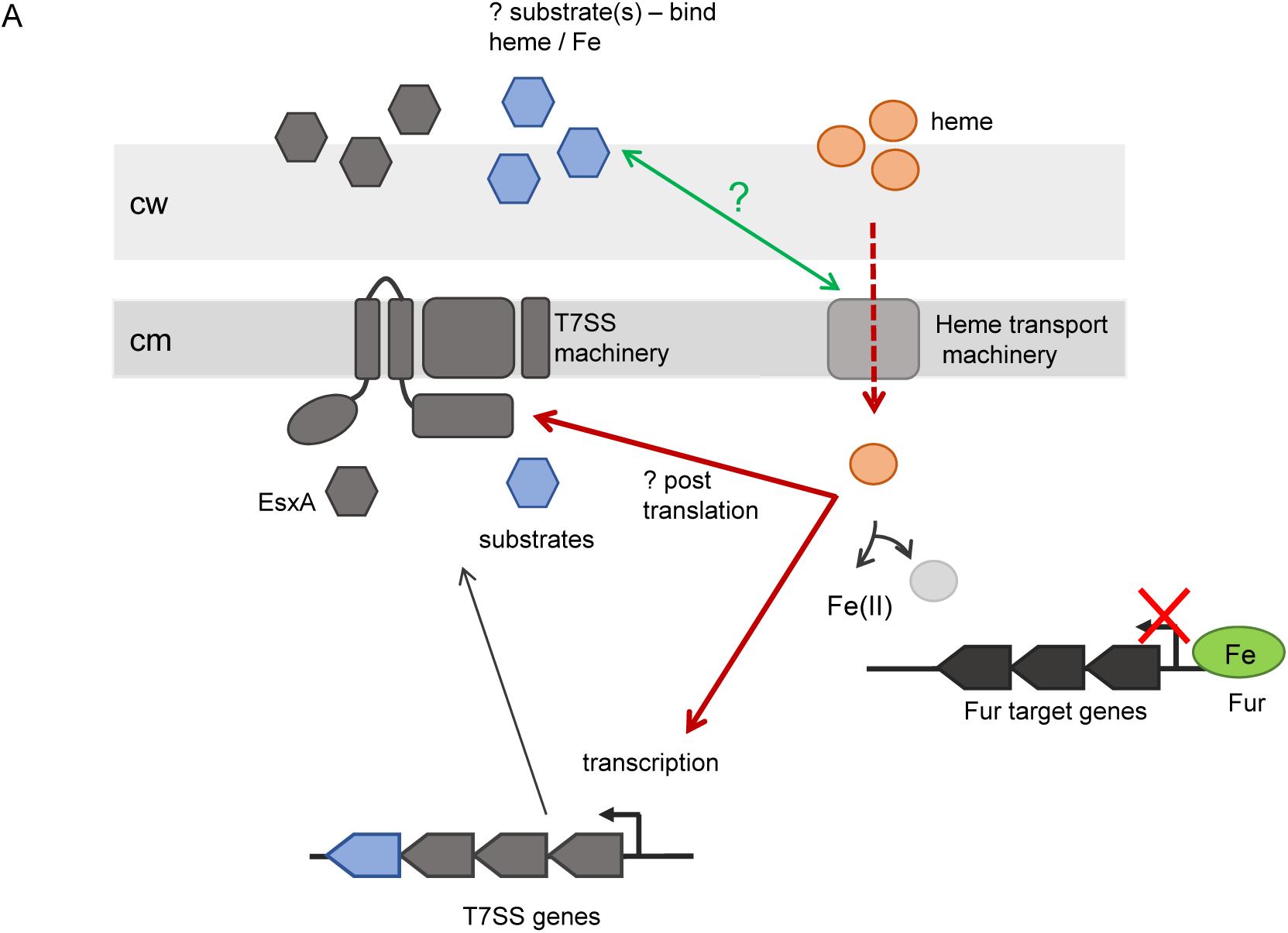
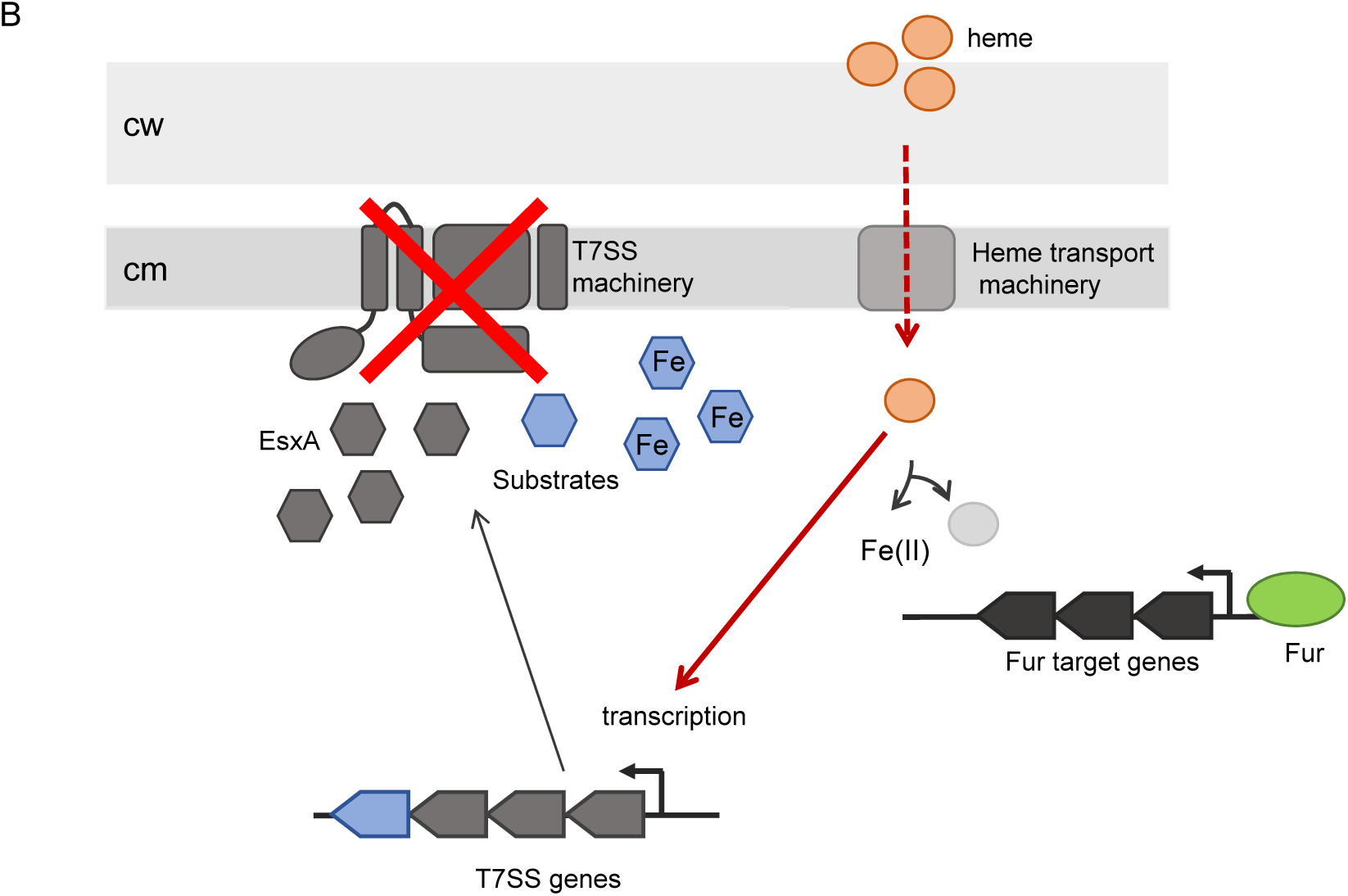
Possible model for the role of the T7SS of *S. aureus* RN6390 strain in iron homoeostasis. (A) The T7SS of *S. aureus* strain RN6390 secretes one or more substrates that can bind heme and/or iron in the extracellular milieu. Increased cytoplasmic levels of heme activate the T7SS at the level of transcription and probably also post-translationally. (B) In the absence of EssC, T7SS substrates accumulate in the cytoplasm, resulting in sequestration of iron in this compartment. This titrates iron away from the iron-binding regulatory protein, Fur, activating transcription of Fur-controlled genes. cm: cytoplasmic membrane, cw: cell wall.

## ACKNOWLEDGEMENTS

This study was supported by the Wellcome Trust (through Investigator Award 10183/Z/15/Z to TP), the Biotechnology and Biological Sciences Research Council and the Medical Research Council (through grants BB/H007571/1 and MR/M011224/1, respectively). The authors would like to thank Professor Ross Fitzgerald (Roslin Institute/University of Edinburgh) for providing us with the 10.1252.X strain, Dr Francesca Short for her assistance with RNA-Seq data analysis, Dr Nicola R. Stanley-Wall for her advice with RNA extraction and Mr Connor Bowen for his assistance in RT-qPCR experiments. Professor Simon Foster (University of Sheffield) is thanked for helpful discussion. The authors declare no conflicts of interest.

## SUPPLEMENTARY FIGURE LEGENDS

**Figure S1.**
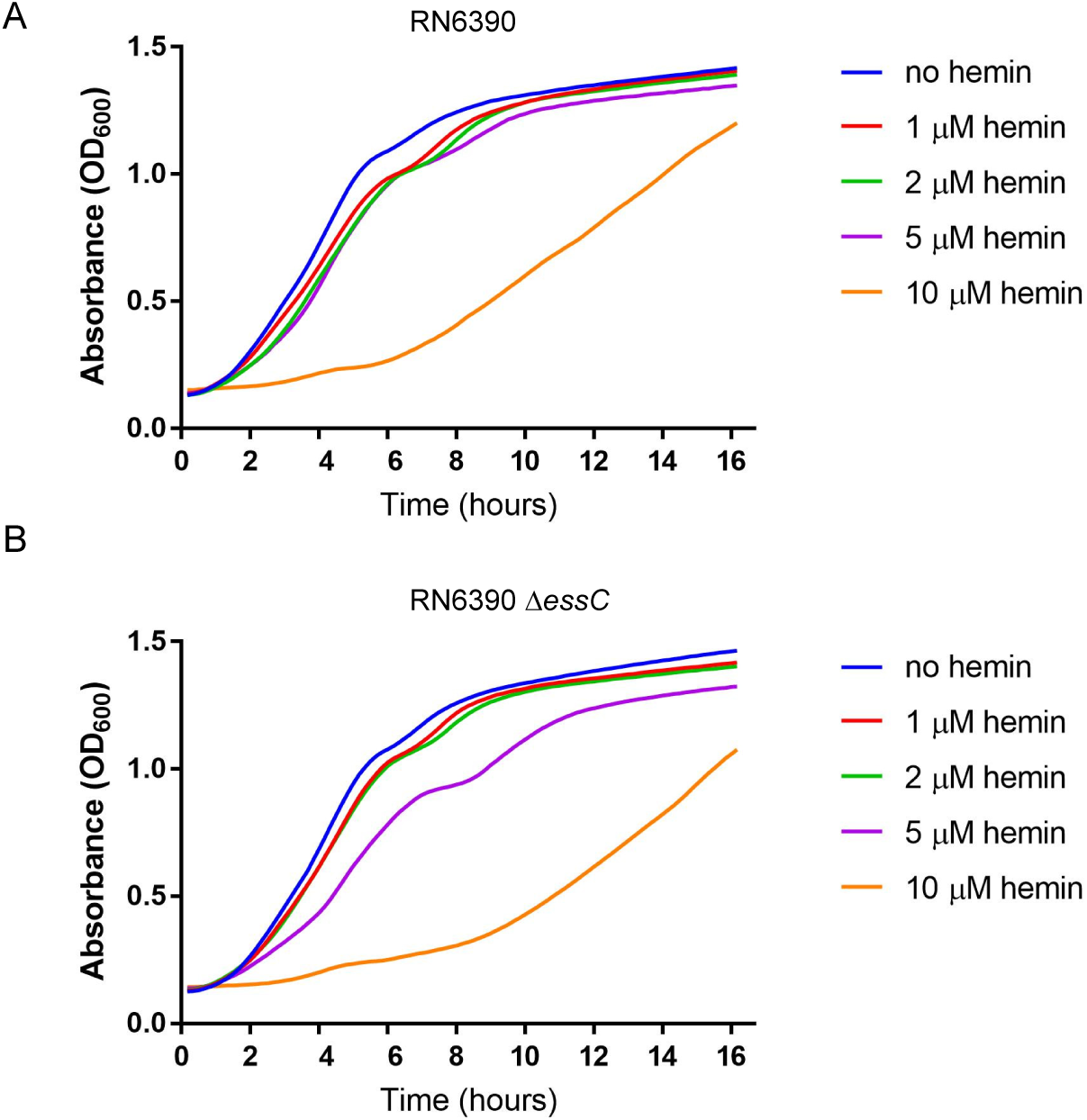
Effect of hemin on the growth of *S. aureus* RN6390. *S. aureus* RN6390 (A) and the isogenic *essC* mutant (B) were grown, with shaking, in TSB medium supplemented with the indicated concentrations of hemin. Growth was monitored over 16 hours in 96-well plates (200 µl volume).

**Figure S2.**
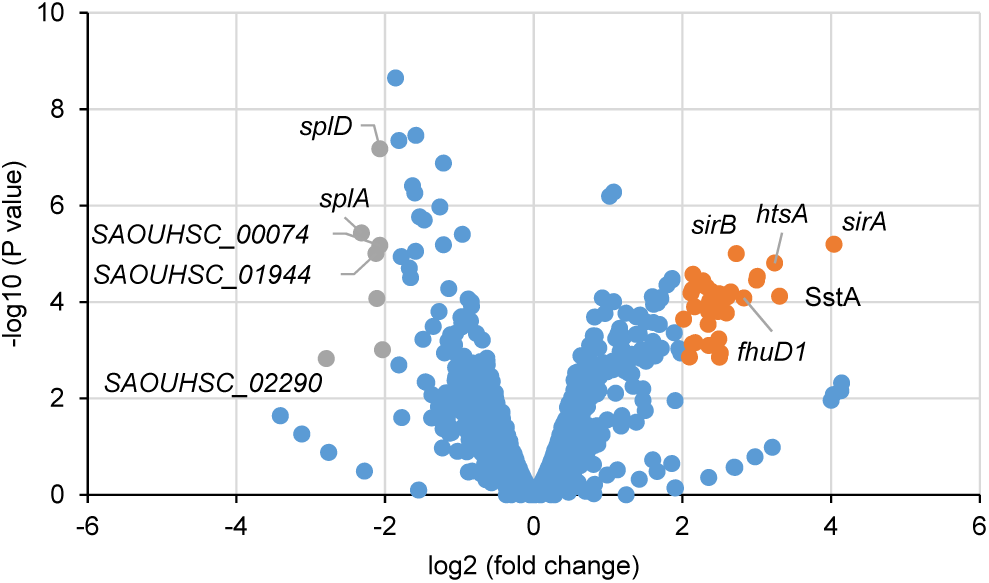
Volcano plot representation of the differentially expressed genes in RN6390 strain compared to the isogenic *essC* mutant. The orange and grey spots represent, respectively, genes that are up- or down-regulated in *essC* mutant relative to the parental strain. Note that the *essC* gene was removed from this analysis.

